## Supplemental Material for "Quantitative Mapping of Keratin Networks in 3D"

**Suppl. Figure 1.** Exemplary representation of major steps in the workflow. All data are in 3D. **(A)** Maximum intensity projection of a small region of interest taken from an airyscan image of fluorescently labelled keratin filaments. (**B)** 'Snakes' prepared by TSOAX segmentation. White dots represent vertices and orange dots represent start and end vertices of 'snakes'. Colored lines distinguish individual 'snakes'. Note that 'snakes' are not well defined since not all 'snakes' have an end at intersections (red arrows). Therefore, the start and end points of snakes are not defined and snakes may span over several intersections. **(C)** In the final segmentation, segments (colored lines) and nodes (red dots) are defined. No intersections of segments exist. Each segment has a start and end point (= node), and segments are connected by nodes. Y-shaped (blue arrow) or X-shaped (red arrow) branchings occur. (**D)** The cinematic rendering shows segments as tubes. Thickness represents mean brightness.

**Suppl. Figure 2. Validation of segmentation. (A)** Maximum intensity projections of airyscan recordings of 10 cells each (left, MDCK; middle, HaCaT; right, RPE. **(B)** The pictures show the xy-position and brightness of segment vertices after 3D-Gaussian blurring. The resulting convolutions mimic those of microscopic imaging. **(C)** The merged images were prepared by overlay of the first and second row above to compare original imaging data with segmentation. **(D)** The picture shows all digital representations prepared from focal planes of airyscan image stacks of single cells. The xy-position of all segments is plotted in their corresponding slice position. Corresponding Video 1 shows an animated version of the data.

**Suppl. Figure 3.** Boxplots of cell volumes estimated from fluorescence image stacks **(A)** and calculated virtual persistence length of segments **(B)** in MDCK, HaCaT and RPE cells.

**Suppl. Figure 4.** Histograms of keratin filament orientation in MDCK, HaCaT, and RPE cells. (**A**) The normalized histograms show the normalized distribution of azimuth (= horizontal orientation) of keratin filament segments in 10 MDCK, HaCaT, and RPE cells, respectively. Since the segments have no defined start and end point, the histograms are restricted to 180 degree. The horizontal line represents the value, if all angles were uniformly distributed. (**B**) The normalized histograms show the elevation (= vertical orientation) of keratin filament segments in 10 MDCK, HaCaT, and RPE cells. The horizontal line delineates the value, if all angles were uniformly distributed.

**Video 1.** Validation of keratin filament segmentation in an MDCK, HaCat and RPE cell. The animation shows the original fluorescence image stacks (left), the segmented data including the associated brightness (middle) and the overlays of both (right).

**Video 2.** Cinematic rendering of the reconstructed keratin filament network of an MDCK cell growing in a confluent monolayer. The tubes represent keratin bundles with different thickness. At the beginning of the video the segments are randomly color-coded.

**Video 3.** Cinematic rendering of the numerical reconstructed keratin network of a HaCaT cell growing in a confluent monolayer. The tubes represent keratin bundles with different thickness. At the beginning of the video the segments are randomly color-coded.

**Video 4.** Cinematic rendering of the numerical reconstructed keratin network of an RPE cell. The tubes represent keratin bundles with different thickness. At the beginning of the video the segments are randomly color-coded.

### 1 Supplemental Figures

#### 1.1 Suppl Fig 01

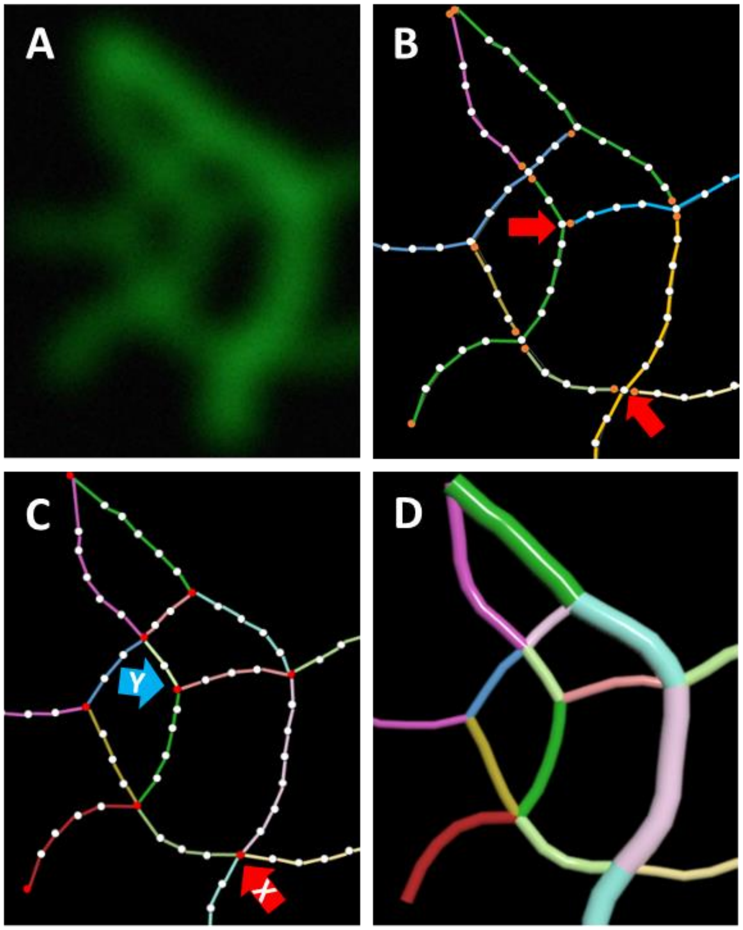

#### 1.2 Suppl Fig 02

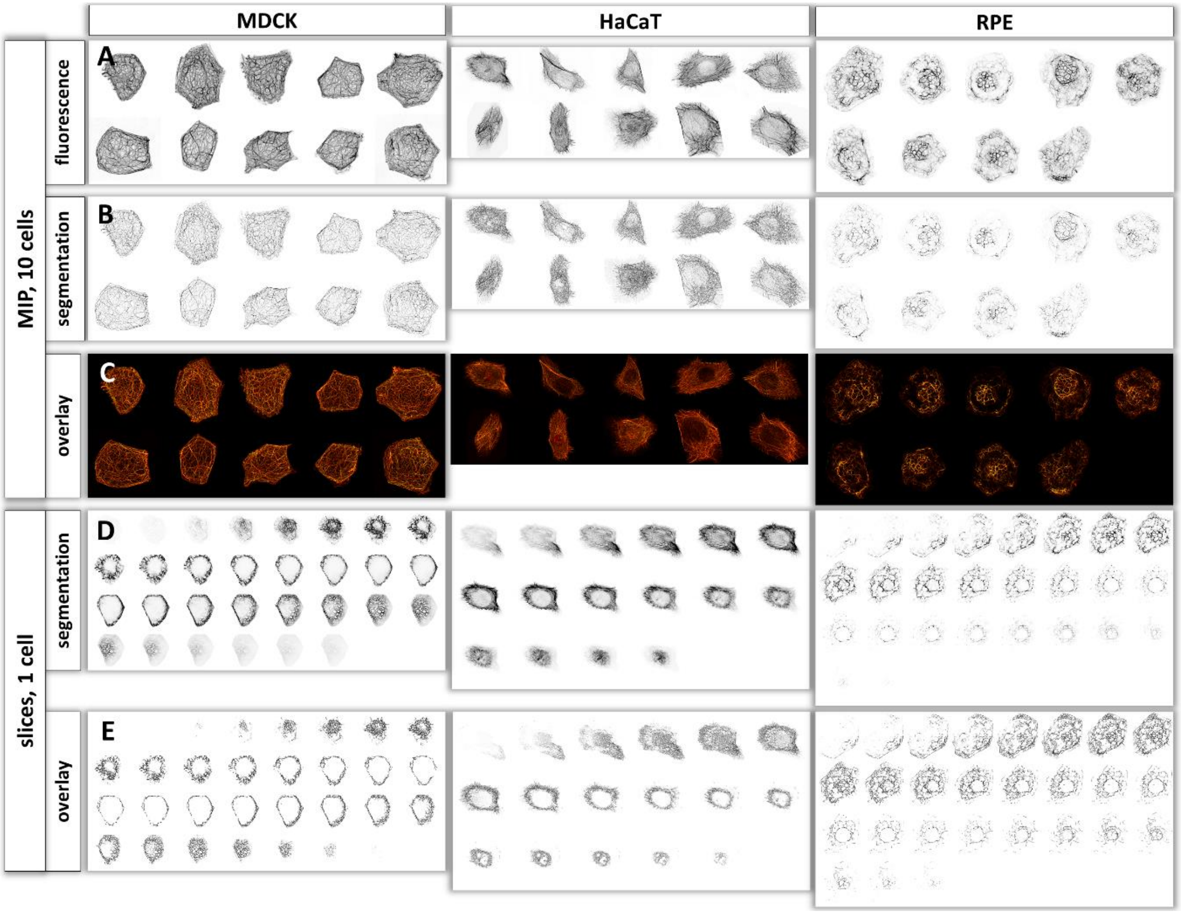

#### 1.3 Suppl Fig 03

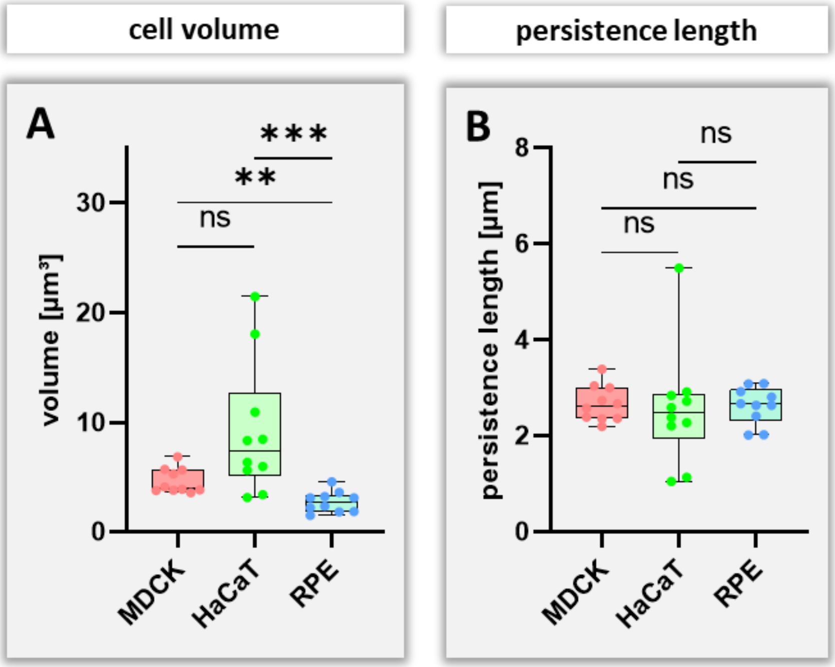

#### 1.4 Suppl Fig 04

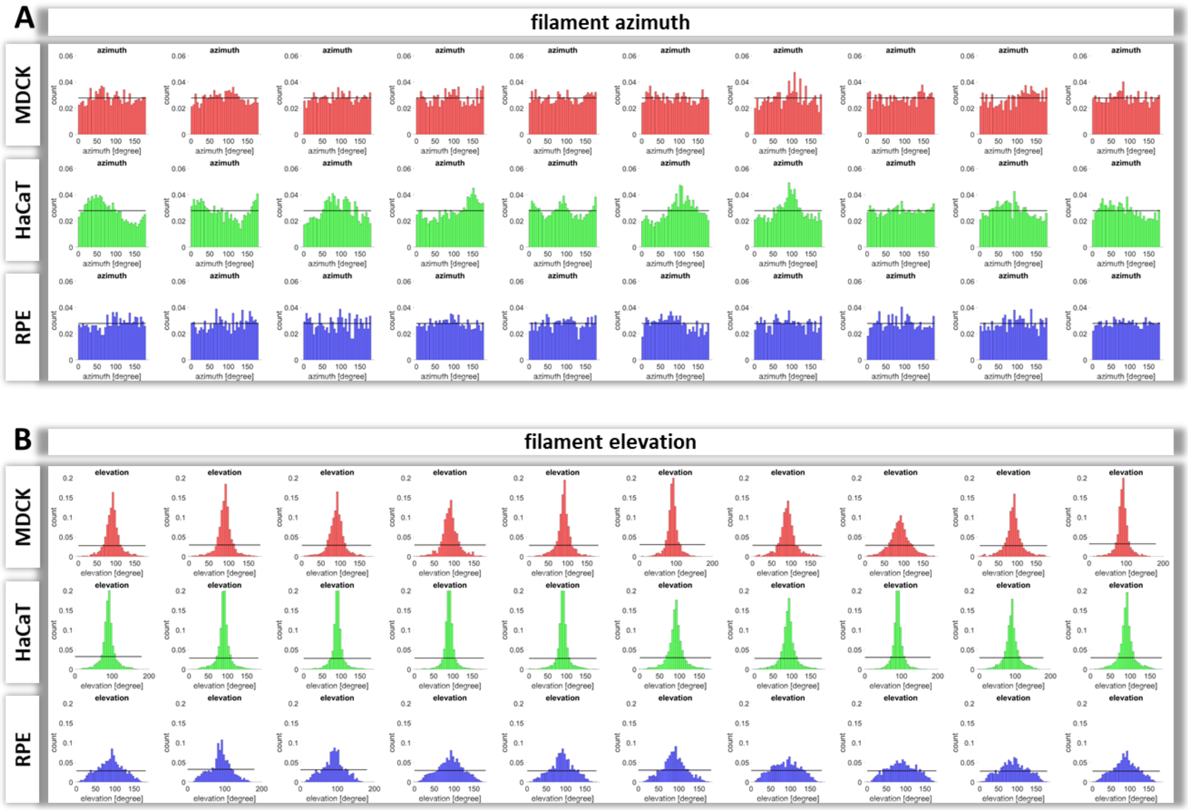

### 2 Videos

#### 2.1 Video 01

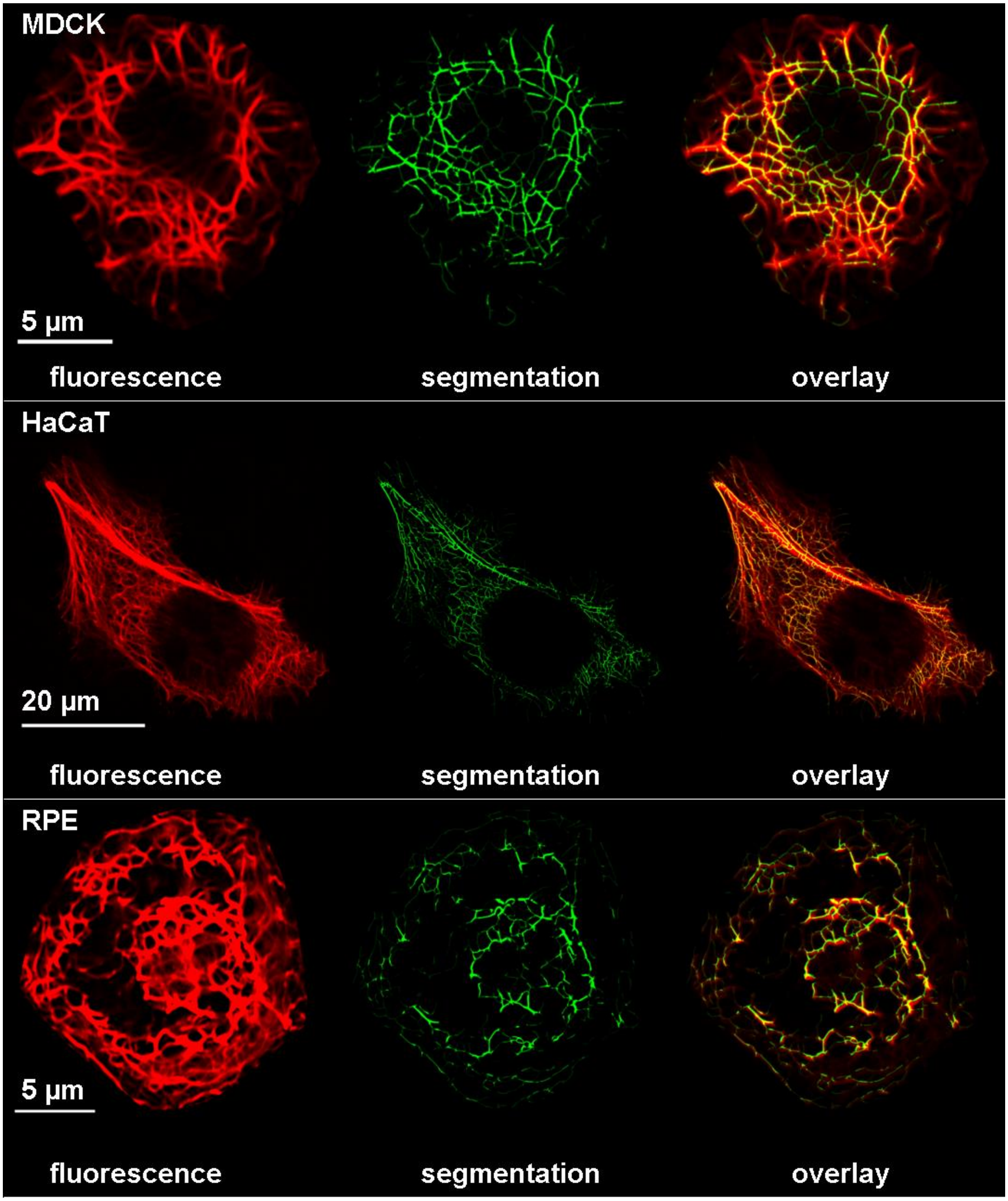

#### 2.2 Video 02

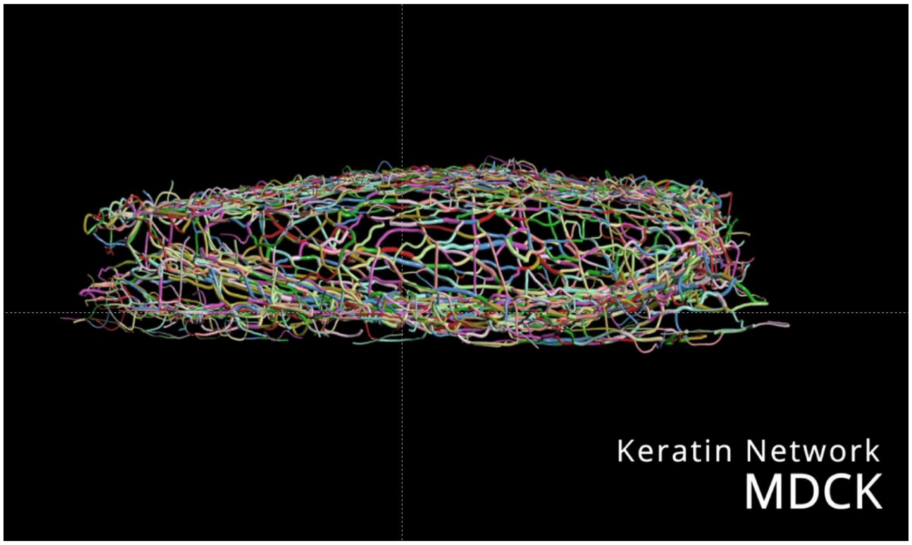


#### 2.3 Video 03

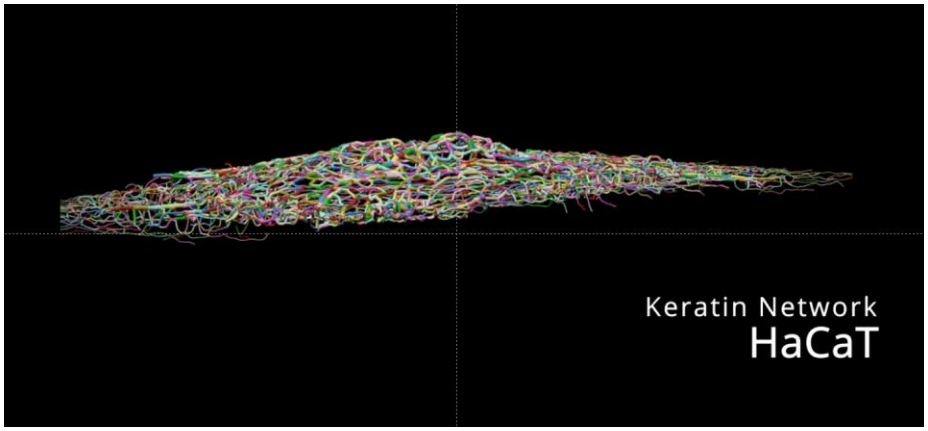


#### 2.4 Video 04

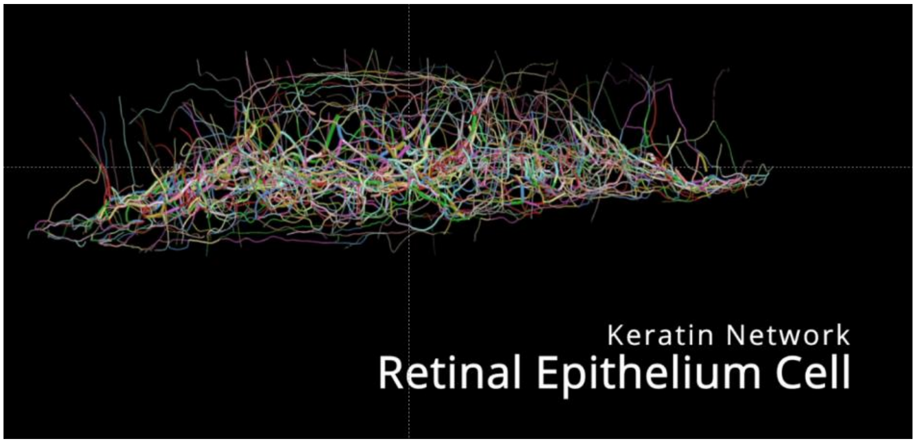
